# Regulatory dynamics distinguishing desiccation tolerance strategies within resurrection grasses

**DOI:** 10.1101/2022.02.16.480747

**Authors:** Brian St. Aubin, Ching Man Wai, Sunil K. Kenchanmane Raju, Chad E. Niederhuth, Robert VanBuren

## Abstract

Desiccation tolerance has evolved recurrently in grasses using two unique strategies to mitigate photooxidative damage under anhydrobiosis. The grass *Oropetium thomaeum* protects and retains chlorophyll, thylakoids, and the photosynthetic apparatus during desiccation (Homoiochlorophyly), while *Eragrostis nindensis* degrades and resynthesizes these components under desiccation and rehydration (Poikilochlorophyly). Here, we surveyed chromatin architecture and gene expression during desiccation in these two closely related species to identify regulatory dynamics underlying the distinct desiccation tolerance strategies in grasses. In both grasses, we observed a strong association between nearby chromatin accessibility and gene expression in desiccated tissues compared to well-watered, reflecting an unusual chromatin stability under anhydrobiosis. Integration of chromatin accessibility (ATACseq) and expression data (RNAseq) revealed a core desiccation response across these two grasses including many genes with binding sites for the core seed development transcription factor ABI5. *O. thomaeum* had a unique set of desiccation induced genes and regulatory elements associated with photoprotection, pigment biosynthesis, and response to high light, reflecting its adaptation of homoiochlorophyly. A tandem array of early light induced proteins (ELIPs) had massive shifts in gene expression and chromatin openness under desiccation in only *O. thomaeum*, and ELIPs acquired a novel desiccation related cis-regulatory motif, reflecting regulatory neofunctionalization during the evolution of desiccation tolerance. Together, our results highlight the complex regulatory and expression dynamics underlying desiccation tolerance in grasses.

## Introduction

Water deficit was the most pervasive challenge faced by charophyte green algae during the colonization of land, and it has been a continual force shaping the evolution and diversification of plants. Plants have evolved a versatile arsenal of strategies to avoid or overcome water limitations. At the extreme, a small group of plants can survive prolonged desiccation for months to years until the return of water. Desiccation tolerance, or the ability to survive atmospheric drying, has been investigated for more than a century (Bewley, 1979; Bristol, 1919). Anhydrobiosis, or ‘life without water’, causes cells to enter a glassy or solid state and puts tremendous stress on all of the macromolecules and organelles. Polysomes are lost during drying, effectively halting protein synthesis, but they are quickly restored during rewetting (Bewley, 1979). Mitochondria and chloroplasts of most species swell and become ill defined during desiccation, and only species able to restore these organelles are associated with recovery from dry conditions (Sherwin and Farrant, 1998; Wellburn and Wellburn, 1976). Biochemical, physiological, and molecular mechanisms underlying desiccation tolerance have been thoroughly described, but comparatively little is known about the regulatory dynamics controlling this trait and what distinguishes desiccation tolerance from more typical drought responses (Gechev et al., 2021).

Desiccation tolerance requires tight coordination of numerous cellular processes and pathways to protect plants from the effects of water deprivation and photooxidative damage under excess light. Various osmoprotectants, heat shock proteins, reactive oxygen species scavengers, changes in membrane lipid composition, and late embryogenesis abundant (LEA) proteins have well-documented roles in desiccation tolerance (Hoekstra et al., 2001). Along with these conserved responses, desiccation tolerant plants utilize two distinct strategies to mitigate photo-oxidative damage. Homoiochlorophyllous species retain and protect their chlorophyll and thylakoids during desiccation whereas poikilochlorophyllous species break down and rebuild chlorophyll, thylakoids, and components of the photosynthetic apparatus under anhydrobiosis using specialized membrane free plastids (desiccoplasts) (Tuba et al., 1994). These divergent strategies are associated with distinct rehydration and photosynthetic kinetics and have utility in different environments.

Vegetative desiccation tolerance is thought to have evolved through rewiring pre-existing pathways, perhaps convergently across independent lineages. Desiccation tolerance mechanisms are broadly conserved between vegetative tissues and seeds, prompting the long-standing hypothesis that desiccation tolerance evolved through rewiring existing seed pathways (Illing et al., 2005; Costa et al., 2017; VanBuren, 2017; VanBuren et al., 2018a). However, recent comparative experiments suggest this connection is more nuanced, as seed pathways also play an important role in typical drought responses of desiccation sensitive species (Pardo et al., 2020). Water deficit responses are broadly regulated by the phytohormone abscisic acid (ABA), and the ABA regulon has a major role in desiccation tolerance (Gaff and Oliver, 2013; Manfre et al., 2009; Shinozaki and Yamaguchi-Shinozaki, 2007) and drought responsive pathways (Daszkowska-Golec, 2016). Several transcription factor families including dehydration-responsive element-binding factor (DREB), basic leucine zipper (bZIP), and NAM-ATAF-CUC2 (NAC) are regulated by ABA and play integral roles in drought and desiccation responses (Nakashima et al., 2014; Takasaki et al., 2015; Wang et al., 2019; Yoshida et al., 2015). The unique and overlapping pathways between drought and desiccation and the regulatory machinery underlying these stress responses remains poorly understood.

The regulation of gene activity based on proximity to *cis*-sequences or entire regions has been a long standing interest of the biological community (Baker, 1968), and has been the source of significant advancement in our understanding of eukaryotic gene regulation (Eissenberg, 1989). High quality analysis of plant chromatin structure and accessibility has been under investigation for several decades (Bowler et al., 2004). Recently, the use of assay for transposase accessible chromatin sequencing (ATAC-seq) has become a powerful tool to reveal binding sites of transcription factors and correlates well with changes in gene expression in various tissues and cell types (Lu et al., 2017). ATAC-seq has even been used to draw correlations between species to identify evolutionarily conserved chromatin regions (Lu et al., 2019). Chromatin dynamics have been profiled under several abiotic stresses including cold, heat, and flooding (Wang et al., 2021; Reynoso et al., 2019; Han et al., 2020; Liang et al., 2021), but little is known about chromatin changes during desiccation. One small scale investigation into cis-element activity found a desiccation-ABARE promoter region to be active in guard cells during desiccation (Smith-Espinoza et al., 2007). However, that study does not give a clear picture of how chromatin is affected by the loss of water, or the link between chromatin changes and gene expression.

Here, we surveyed changes in chromatin architecture, gene expression, and regulatory dynamics under desiccation in two related resurrection grasses with contrasting photoprotective strategies. We collected parallel datasets from the homoiochlorophyllous (chlorophyll retaining) model resurrection grass *Oropetium thomaeum* and the poikilochlorophyllous (chlorophyll degrading) grass *Eragrostis nindensis*. These species are found in the Chloridoideae subfamily of grasses, a group of diverse, stress tolerant species that includes the important underutilized crops tef (*Eragrostis tef)* and finger millet (*Eleusine coracana*). Oropetium has the smallest diploid sequenced genome among grasses (∼250 Mbp) (VanBuren et al., 2018b). Studies using Oropetium as a model for desiccation have revealed metabolic characteristics common among organisms capable of anhydrobiosis such as changes in sugar metabolism and these changes are specific to desiccation compared to other stresses (Bartels and Mattar, 2002). Previous studies in *O. thomaeum* showed an induction of gene families such as ELIPs and LEAs during desiccation, and we sought to test if there were similarities in chromatin accessibility near these genes (VanBuren et al., 2017). Using this comparative system, we show that significant shifts in chromatin architecture are associated with desiccation and photoprotective responses in resurrection grasses. We identified a core set of genes, open chromatin peaks, and cis-regulatory elements that underly conserved desiccation tolernace responses across these two grasses. We also found distinct regulatory dynamics that are associated with the different strategies of retaining or degrading the photosynthetic apparatus under anhydrobiosis.

## Results

### Dynamic patterns of chromatin accessibility under desiccation

We surveyed the changes in chromatin architecture and gene expression underlying desiccation tolerance using a combination of ATAC-seq and RNAseq in *O. thomaeum* and *E. nindensis*. Mature plants were slowly desiccated over a period of ten days without watering, and samples for RNAseq and ATACseq were collected in parallel from leaves of desiccated and well-watered plants (Figure 1). Expression dynamics were profiled using RNAseq, and quality trimmed reads were pseudo-aligned to the *O. thomeaum* and *E. nindensis* gene models using Kallisto (Bray et al., 2016). Differentially expressed genes were identified using Sleuth (Pimentel et al., 2017). In *O. thomaeum*, comparison of RNA-seq data from well-watered and desiccated plants revealed 5,187 genes with higher expression under desiccation and 5,354 genes with lower expression compared to well-watered (q < 0.05). Similar timepoints in *E. nindensis* had 8,071 genes with higher expression under desiccation and 8,971 genes with lower expression compared to well-watered. This represents roughly one third of the genes in the *O. thomaeum* genome and almost half of the genes with detectable expression. In *E. nindensis*, the differentially expressed genes represent about 15% of all genes and 30% of the genes with detectable expression. This highlights the complex transcriptional reprogramming required for the successful deployment of desiccation tolerance. The *O. thomaeum* genes with the highest expression in desiccation include dehydrins (Ot_Chr8_n39154), early light induced proteins (ELIPs; Ot_Chr8_39125, Ot_Chr8_n38842), and other proteins with well-characterized roles in desiccation and abiotic stress responses (Supplemental Table 1). Genes with the highest differential expression in *E. nindensis* desiccated plants included two late embryogenesis related proteins (En_0084954, En_0002557), an aldehyde dehydrogenase (En_0060658), and a glucosyl transferase (En_0017075; Supplemental Table 2).

**Figure 1.**
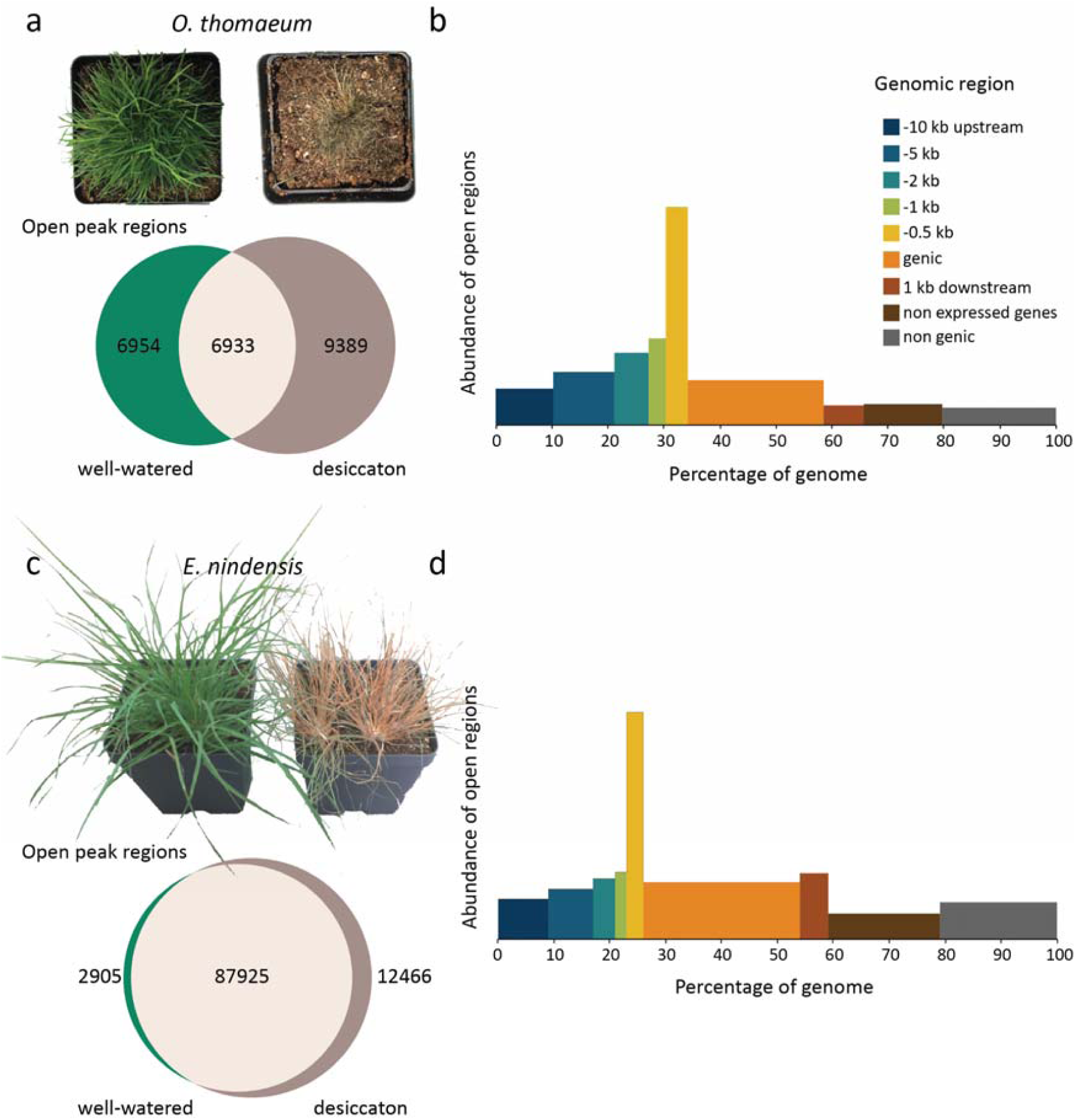
Chromatin architecture changes underlying desiccation responses in resurrection grasses. (a) Overlapping (beige) open chromatin regions between well-watered (green) and desiccated (dark grey) *O. thomaeum* plants. Representative well-watered (left) and fully desiccated (right) *O. thomaeum* plants are shown. (b) Landscape of open chromatin regions across the *O. thomaeum* genome. The abundance of open chromatin regions is shown on the Y axis and percentage of the genome corresponding to upstream, genic, downstream, intergenic, and genes with no detectable expression is shown on the X axis. (c) Overlapping (beige) open chromatin regions between well-watered (green) and desiccated (dark grey) *E. nindensis* plants. Representative well-watered (left) and fully desiccated (right) *E. nindensis* plants are shown. (b) Landscape of open chromatin regions across the *E. nindensis* genome.

Using ATAC-seq, we were able to quantify chromatin openness in desiccation and well-watered conditions for the same tissue analyzed by RNA-seq. Nuclei were isolated from flash frozen samples and ∼50,000 nuclei were used to construct ATACseq libraries with two biological replicates per treatment. Trimmed reads were aligned to the *O. thomeaum* V2 or *E. nindensis* V2 genomes using bowtie2 (Langmead and Salzberg, 2012) with the overall alignment rates varying from 85% to 97% across all samples. Peaks of open chromatin were called using Genrich (Gaspar) and visual inspection of the aligned ATACseq reads in a genome browser(Supplemental Figure 1 and 2). Peaks are strong and consistent across replicates and treatments after removal of regions likely to be chloroplast contamination, indicating the libraries are high quality. Differential peak calling with DiffBind (Stark et al., 2011) provided a Fraction of Reads in Peaks (FRiP) of between 0.27 and 0.49. The well-watered samples all have lower FRiP than the desiccated samples, however fewer open regions were detected in desiccated samples with 18,112 vs. 28,538 in *O. thomaeum*, and 59,514 vs. 107,593 in *E. nindensis*.

In total, we identified 30,134 accessible chromatin regions (ACRs) in *O*.*thomaeum* and 121,999 ACRs in *E. nindensis* occupying 4.6 % and 7.0 % of the genome respectively (Figure 1). Roughly 85% and 82% of all ACRs were near expressed genes in *O. thomaeum* and *E. nindensis*, respectively. For both species, the 0-500 bp bin upstream of the transcription start site (TSS) has an over-representation of ACRs based on the relative proportion of the genome occupied by this range (Figure 1b, d). We also observed that ACR density decreases with distance from the TSS. The *O. thomaeum* and *E. nindensis* genomes are highly compact compared to other grassses (VanBuren et al., 2015, 2020; Pardo et al., 2020), and expressed genes and their flanking regions (10kb upstream to 1kb downstream) cover 65.8% and 59.4% of the grass genomes respectively. Not all ACRs were found near canonical genes. Of the open regions, 7.2% and 14.2% are near non-expressed genes and 7.5% and 21.5% of the regions were not near any genes. We inspected the sequences of the regions not near genes and found several motifs present in multiple regions. This could imply an evolutionary significance for these regions that will need to be investigated at a future time.

### Chromatin architecture is tightly linked to the regulation of desiccation responsive genes

Since chromatin openness is believed to impact the expression level of nearby genes, we investigated the relationship between chromatin openness and gene expression changes due to desiccation using two different methods. Expressed genes typically have open chromatin near the TSS (Figure 1b, d), and we used deepTools (Ramírez et al., 2016) to aggregate the genomic regions near differentially expressed genes. On average, we found small chromatin regions near TSSs which often extend beyond 1kb upstream of the gene and slightly into the gene body (50-100bp) that have significantly more openness when the gene has higher expression (Figure 2). This effect was most pronounced for genes with higher expression in desiccation in both *O. thomaeum* and *E. nindensis*. Genes with upregulated expression in well watered tissue had a distinct peak in openness near the TSS, but this was less pronounced than desiccation induced genes. We then looked at differentially expressed genes that had a differentially accessible region in each bin and plotted the best-fit line for each combination. The R^2^ value for that line-fit drops off inside the 5000 to 2000bp region and is highest for the 0 to 500bp region upstream of the TSS (Supplemental Figure 3).

**Figure 2.**
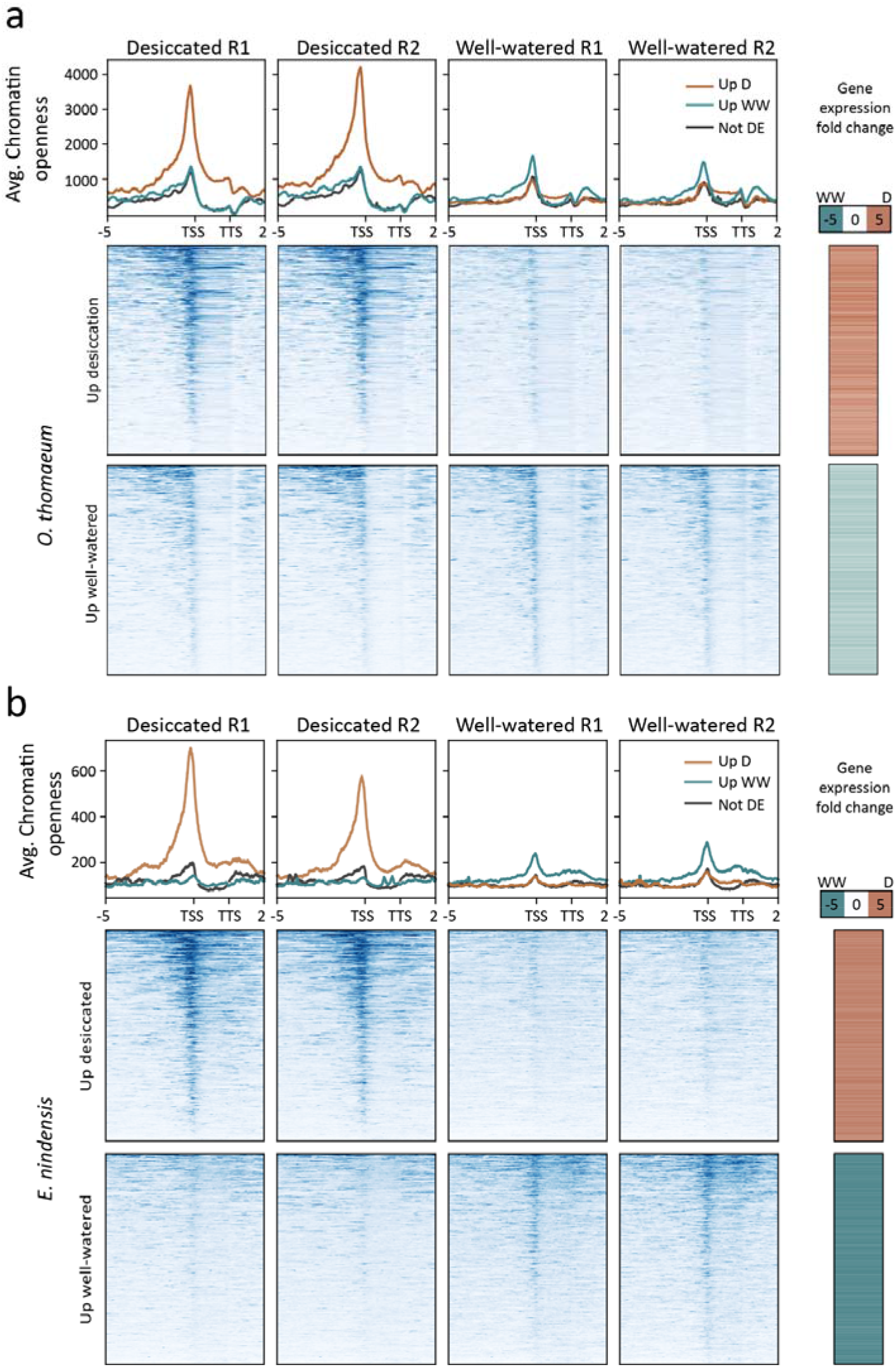
Chromatin architecture and gene expression associations under desiccation. The average chromatin openness is plotted for genes upregulated under desiccation (orange) or upregulated under well-watered (blue) for the two replicates in each sample for *O. thomaeum* (a) and *E. nindensis* (b). The log2 fold change of expression in desiccated / well-watered is plotted on the right for each gene. Openness is plotted 5 kbp upstream to the TSS, on the gene is between TSS and TTS,and TTS to 2 kbp downstream. TSS: Transcriptional start site, TTS: transcription termination site.

**Figure 3.**
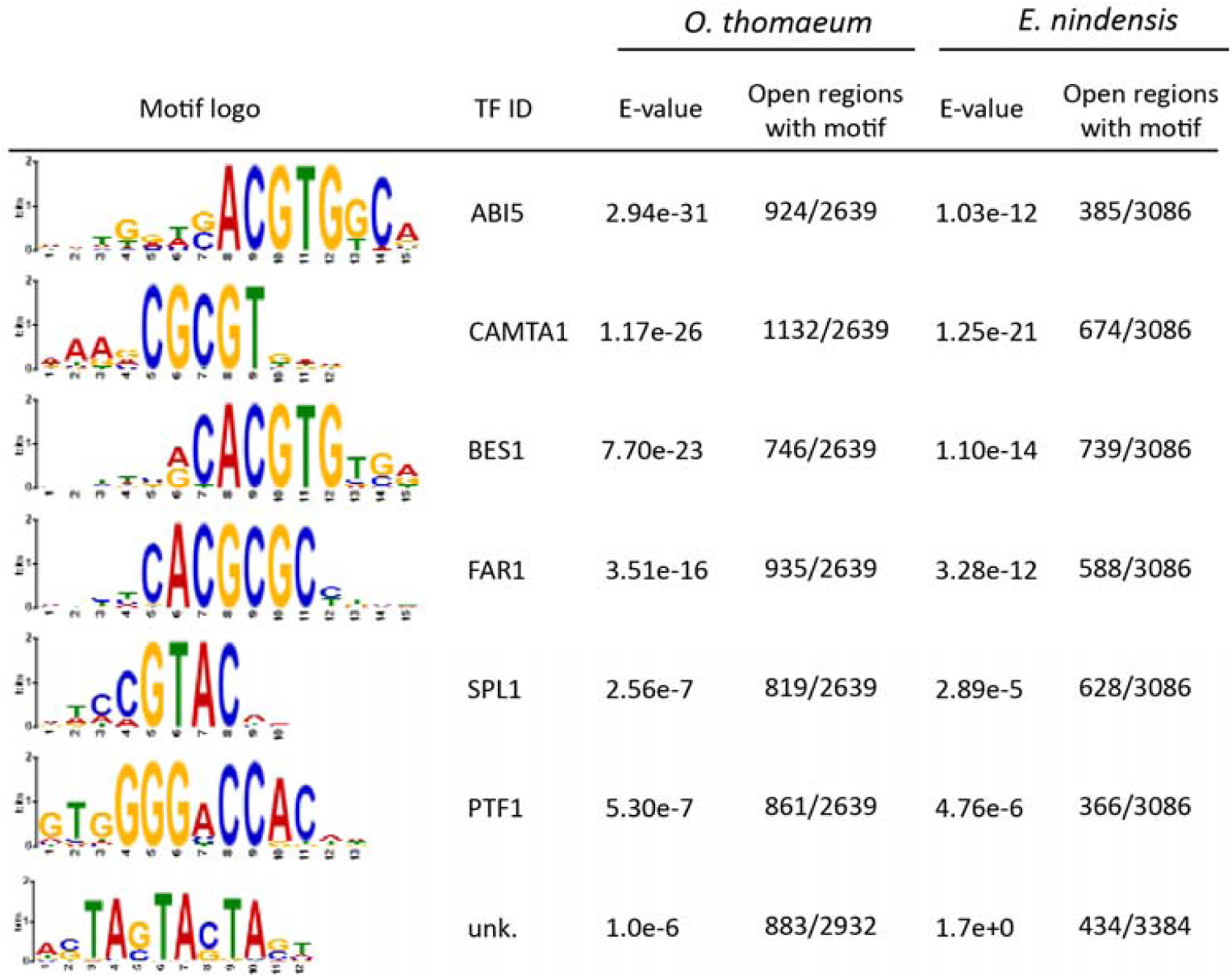
Enriched regulatory motifs associated with desiccation responsive genes. Enriched *cis*-regulatory element motifs in open chromatin regions of genes upregulated under desiccation in both grasses are shown. The last logo (TF ID: unknown) was obtained from a de novo search for motifs. The motif logo, most closely associated transcription factor, statistical significance (E-value), and number of open regions with each motif is shown for *O. thomaeum* (left) and *E. nindensis* (right).

To better understand gene expression during desiccation and the impacts that open chromatin may have on expression dynamics, we split the differentially expressed genes into four categories depending on the direction of differential gene expression and chromatin openness. There are 18,344 expressed genes in *O. thomaeum*, and 15,880 (86.6 %) have an ACR nearby. Of the 10,542 differentially expressed genes, we found 9,213 (87.4%) to have an ACR nearby. Genes with higher expression in one treatment were more likely to have a chromatin region that was more open nearby. Well-watered samples had 2,330 cases, and desiccated samples had 3,314 cases where a higher expressed gene was near a more open ACRs. Whereas there were only 424 and 1,307 cases of higher expression in desiccation with more open ACRs in well-watered or vise-versa. In *E. nindnesis* there were 65,205 expressed genes with 41,756 (64.0%) having an ACR nearby. We found 17,043 differentially expressed genes with 13,894 (81.5 %) having an ACR nearby. The well-watered samples had 459 cases and desiccated samples had 3,609 cases where a higher expressed gene was near a more open ACR. The genes with higher expression in desiccation but a more open region when well-watered included 47 cases, and there were 786 with the reverse.

To enable detailed comparisons across species, we utilized a set of syntenic orthologs between the two species for expression comparisons (Pardo et al., 2020). Of the 13,417 differentially expressed syntenic genes in *E. nindensis*, 49.8% (6685) have a syntelog in *O. thomaeum* that is also differentially expressed in the same direction (Figure 4a). Whereas 6.9% (932) of the differentially expressed syntenic genes in *E. nindensis* had opposing expression patterns in *O. thomaeum*. In *O. thomaeum*, we found a similar 48.2% (4,460 of 9,252) of differentially expressed syntelogs with the same expression pattern in *E. nindensis*, and 7.9% (735) with the opposite pattern. We also analyzed sets of differentially expressed genes that also contained differential ACRs nearby. Genes with overlapping expression and accessibility dynamics are associated with the same GO-terms with a few differences noted below.

**Figure 4.**
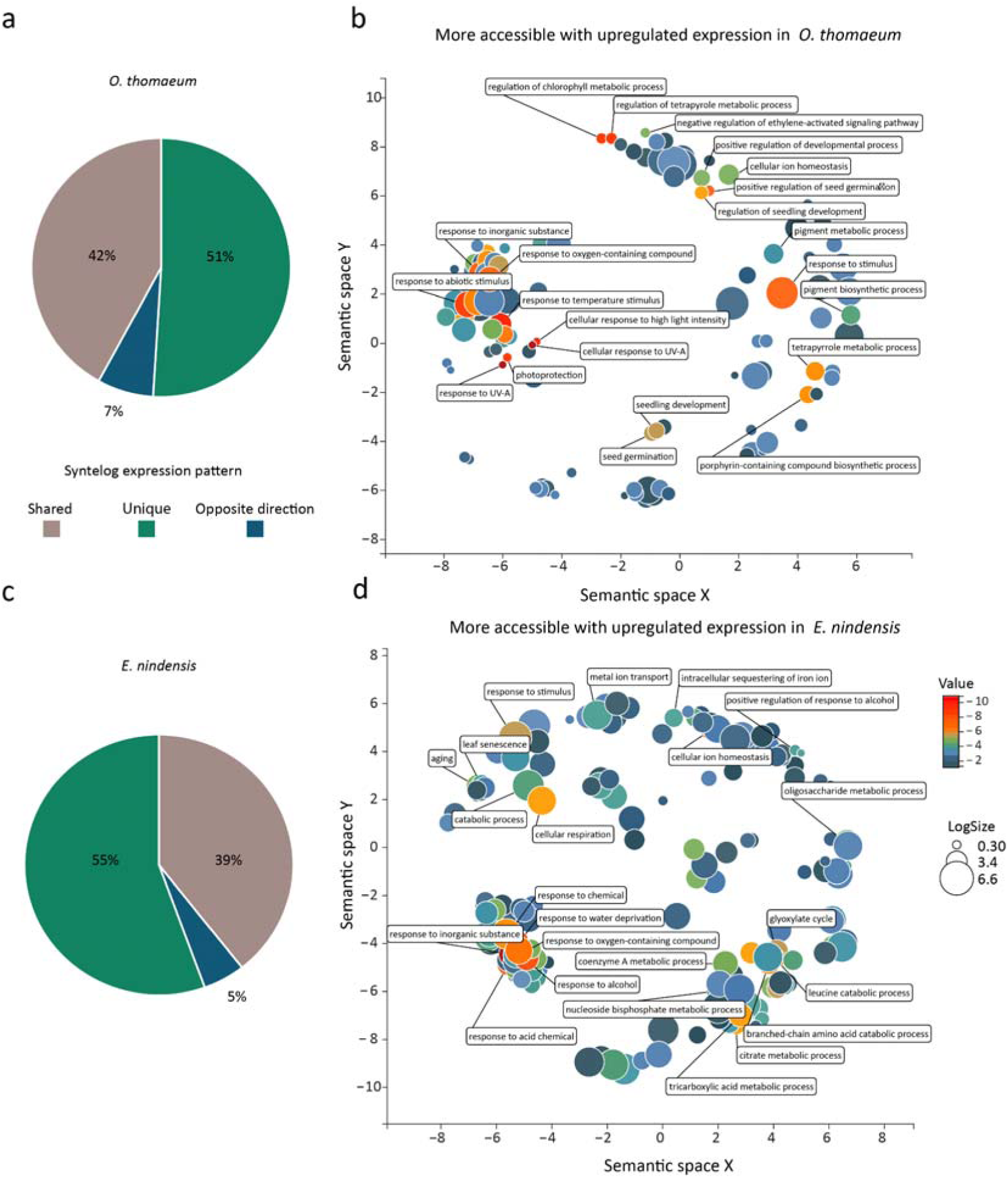
Contrasting enrichment patterns of desiccation-associated genes in *O. thomaeum* and *E. nindensis*. (a) The proportion of syntelogs with similar, unique, or opposing expression patterns is plotted for *O. thomaeum* (left) and *E. nindensis* (right). Enriched GO terms for genes that have more open chromatin and higher expression under desiccation are plotted for *O. thomaeum* (b) and *E. nindensis* (c). GO terms are transformed using Multidimensional Scaling to reduce dimensionality and terms are grouped by semantic similarities. GO terms with previously characterized roles in desiccation and photoprotection responses are highlighted. The color of the circles represent significance and size represents the number of genes in that group.

We searched for patterns of functional enrichment of syntelogs with similar chromatin and expression dynamics that could indicate conserved underlying desiccation responses in grasses. Enriched GO-terms of upregulated syntelogs in both species include response to water deprivation and other abiotic stresses, fatty acid biosynthesis, golgi organization, proteasomes, and respiration (Supplemental Table 9). GO-terms associated with syntelogs that are similarly down-regulated in desiccation are related to photosynthesis, core and secondary metabolism, cell wall processes and hormone metabolism (Supplemental Table 9).

### Cis-regulatory elements associated with desication tolerance

Using these datasets of open chromatin regions and gene expression, we identified putative *cis*-regulatory elements that are involved in the regulation of desiccation responses. We extracted sequences within differential ACRs near expressed genes in *O. thomaeum* and *E. nindensis* and searched for enriched motifs within each set. We utilized the Simple Enrichment Analysis (SEA) algorithm from Multiple Em for Motif Elicitation (MEME) to identify motifs with similar sequences to those that have been previously investigated (Bailey and Grant, 2021). We also identified novel motifs in the open regions using the Sensitive, Thorough, Rapid, Enriched Motif Elicitation (STREAM) algorithm from MEME. The motifs found using STREAM with the smallest *p*-values were sent to TOMTOM to search against the Arabidopsis transcription factor motif database (Bailey, 2020; Bailey et al., 2009; Gupta et al., 2007; O’Malley et al., 2016). We identified 327 and 251 enriched (E-value ≤ 10) motifs from regions that were more open in desiccation and close to genes with higher expression under desiccation in *O. thomaeum* and *E. nindensis*, respectively (supplemental dataset 1). We found 180 and 46 enriched motifs in more accessible regions near upregulated genes in well-watered *O. thomaeum* and *E. nindensis* leaves, respectively. 68 and 17 enriched motifs were identified from regions that were more open in well-watered and close to genes with higher expression in desiccation in *O. thomaeum* and *E. nindensis*. 274 and 232 enriched motifs from regions that were more open in desiccation and close to genes with higher expression in well-watered in *O. thomaeum* and *E. nindensis* were identified respectively. Enriched motifs under desiccation in *O. thomaeum* and *E. nindensis* are similar (Figure 3), and resemble motifs for transcription factors associated with response to water stress (CAMTA1, ABF2), heat stress (SPL1), seedling development (AREB3, ABI5), and UV protection/light signaling (BES1, PTF1, FAR1) (Kim et al., 2002) (Figure 3). STREAM identified 15 novel motifs from the *O. thomaeum* desiccation samples including a highly enriched motif with no homology in Arabidopsis, Mouse, or Fruitfly datasets (TA(G/C)TA(G/C)TA; 883 of 2932 sites; E-value of 1.0E-06) (Bailey, 2020). This motif was also found in 434 of 3384 sites in *E. nindensis*, however it was not significantly enriched (E-value of 1.7E+0). This motif may play a central role in anhydrobiosis related processes in grasses.

### Regulatory dynamics distinguishing photoprotective strategies under desiccation

*O. thomaeum* and *E. nindensis* utilize different strategies to mitigate photooxidative damage under anhydrobiosis. *O. thomaeum* retains and protects while *E. nindensis* degrades and resynthesizes chlorophyll, thylakoid membranes, and components of the photosynthetic apparatus during desiccation and rehydration cycles. We searched for unique ACRs, expression dynamics, and putative *cis-*regulatory elements that distinguish the distinct desiccation tolerance strategeis in these two grasses. The sets of syntenic orthologs and ACRs between *O. thomaeum* and *E. nindensis* were used for comparative analyses.

Syntelogs or species-specific genes that are uniquely upregulated in *O. thomaeum* had functions related to response to heat and high light, photoprotection, protein localization to chloroplast, regulation of seed germination, and regulation of cell cycle (Supplemental table 9). Upregulated genes with more open ACR in *O. thomaeum* have similar enrichment patterns including regulation of chlorophyll biosynthetic process and cellular response to blue light (Supplemental table 8). Uniquely downregulated genes under desiccation in *O. thomaeum* were enriched in GO terms associated with pollen germination, translation, vitamin B6 biosynthesis, and small GTPase mediated signal transduction (Supplemental Table 9).

Genes that are uniquely upregulated in *E. nindensis* under desiccation include enriched GO terms related to endocytic recycling, glyoxylate cycle, hormone processes, response to hypoxia, and salt stress responses (Supplemental Table 9). Uniquely downregulated genes under desiccation in *E. nindensis* are enriched in GO terms associated with regulation of photosynthetic acclimation, regulation of cell death, response to jasmonic acid, auxin efflux, terpenoid transport and catabolic process, and pectin catabolic process (Supplemental Table 9).

The evolution of desiccation tolerance is associated with massive duplication of early light induced proteins (ELIPs), which play a central role in photoprotection under anhydrobiosis (VanBuren et al., 2019). All sequenced resurrection plants contain large tandem arrays of ELIPs including *O. thomaeum* (20 ELIPs) and *E. nindensis* (27), but chlorophyll retaining species typically have more ELIPs when accounting for ploidy (VanBuren et al., 2019). The 20 ELIPs in *O. thomaeum* are among the most highly expressed genes under desiccation, with on average > 20 fold higher expression compared to well-watered (Figure 5a). The ELIPs have no significant open chromatin peaks under well-watered conditions, but have massive peaks of open chromatin upstream of the TSS under desiccation. All but four of the ELIPs in *O. thomaeum* are found in a single tandem array with high sequence homology. Interestingly, some genes within this array have unique peaks in different positions upstream of the TSS, likely corresponding to changes in non-coding sequences during their duplication. In the conserved peak upstream of each ELIP TSS, we found a highly enriched *cis-*regulatroy motif for the central ABA responsive drought transcription factor AP2/DREB (Figure 5b). This motif is notably absent from the two Arabidopsis ELIPs, and may be related to the evolution of desiccation tolerance. Several other enriched cis-regulatory motifs were identified in a subset of peaks near ELIPs related to light responses (FAR1), ABA signaling (STZ, C2H2), and development (MYB88) (Figure 5b). The 27 ELIPs in *E. nindensis* have a diverse response to desiccation. We found two ELIPs to have no RNA expression, 14 with no differential expression between treatments, seven down regulated under desiccation, and four up regulated under desiccation (Supplemental Figure 6). Two of the 23 open chromatin regions near *E. nindensis* ELIPs were significantly more open under desiccation and near genes with higher expression in desiccation, but the remainder showed no changes. These two differentially-open regions contain sequences similar to the AP2/DREB motifs found near the *O*.*thomaeum* ELIPs. However, a search for enriched motifs among open regions near all *E*.*nindensis* ELIPs did not result in an overlap with motifs found in *O. thomaeum* ELIP open regions. Together, this highlights the important role that ELIPs play in chlorophyll retaining species and a possible regulatory neofunctionalization during the evolution of desiccation tolerance.

**Figure 5.**
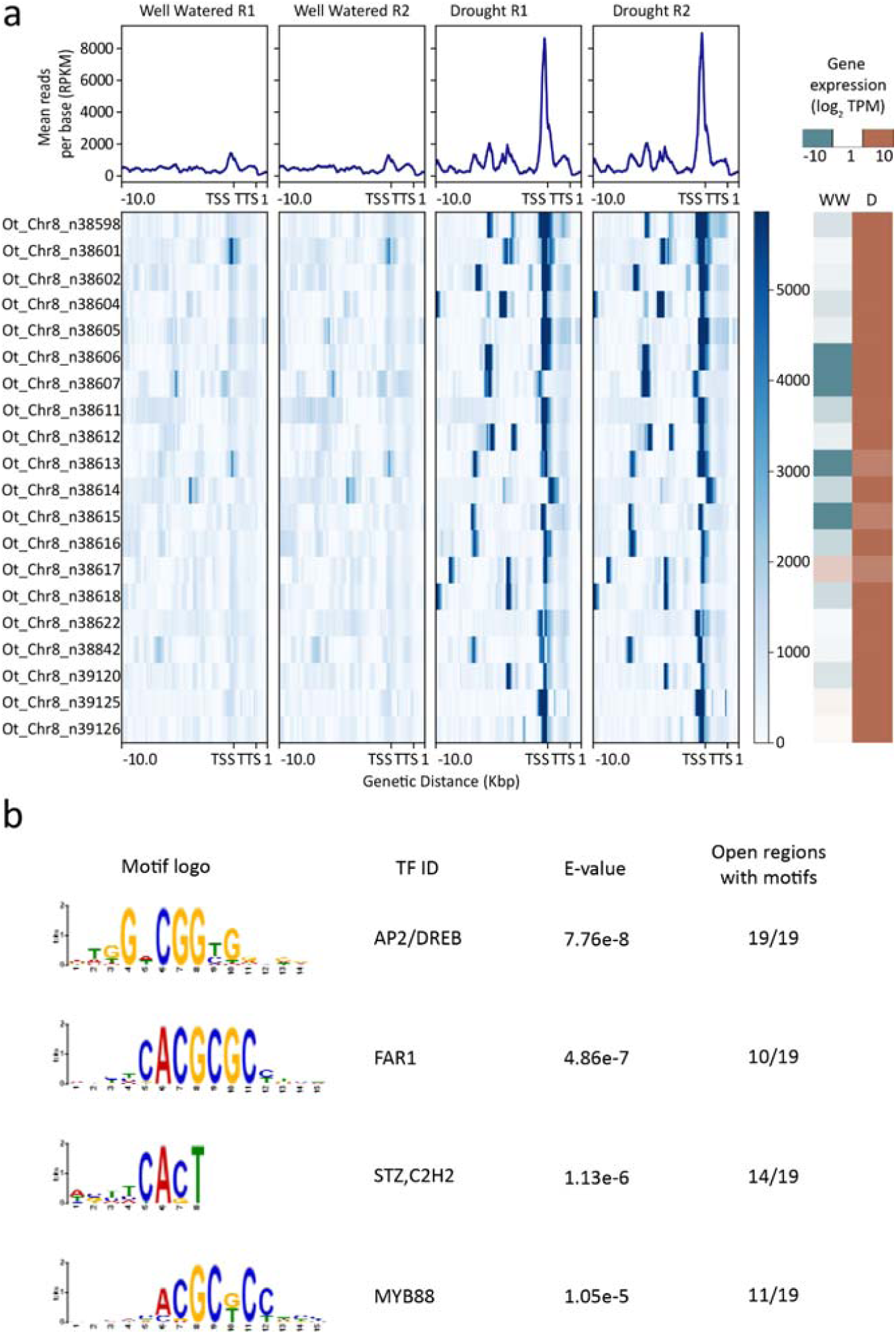
Regulatory dynamics of early light induced proteins (ELIPs) under desiccation. (a) Chromatin architecture and expression dynamics of ELIPs in well-watered and desiccated *O. thomaeum* samples. The mean mapped read depth of ATACseq reads (in RPKM) is plotted for 10 kb upstream to 1 kb downstream regions for each of the ELIPs in the *O. thomaeum* genome. Log2 transformed RNA expression (in TPM) for each ELIP is shown on the right under well-watered and desiccated conditions. (b) Enriched putative *cis*-element motifs in open chromatin regions upstream of ELIP TSSs in *O. thomaeum*. The motif logo, associated transcription factor (TF ID), e-value, and number of open regions with each motif are shown.

## Discussion

Desiccation tolerance has evolved recurrently across the tree of life as a common adaptation to survive anhydrobiosis in water-limited environments. Tolerance is not a static or monolithic trait, and it has been repeatedly gained, lost, or modified across plant evolution. Here, we surveyed changes in chromatin architecture and gene expression in two related resurrection grasses that utilize different photoprotective strategies under anhydrobiosis. *O. thomaeum* and *E. nindensis* have conserved gene content and relatively small monoploid genome sizes (250 and 500 Mb, respectively), enabling detailed comparative genomic analyses and associations between putative cis-regulatory elements and gene function. Open chromatin regions were found in largely genic regions, and desiccation induced shifts in chromatin accessibility for 7.4 Mb (3.1%), and 16.9Mb (1.7%) of the *O. thomaeum* and *E. nindensi*s genomes respectively. These shifts in accessibility are highly correlated with gene expression dynamics, and these high-quality datasets allowed us to explore the unique biophysical constraints of anhydrobiosis and the regulatory evolution of this complex adaptation.

### Chromatin architecture and anhydrobiosis

Chromatin dynamics have not been described for desiccated vegetative tissues, but chromatin is highly condensed and compacted in desiccated seeds (van Zanten et al., 2011). Differential chromatin accessibility is established in developing seeds and gene level modifications enable rapid transcription for germination related processes upon hydration (Fransz and de Jong, 2011). We observed a high signal-to-noise ratio (FRiP) of the open chromatin regions in desiccated leaf tissue stemming from a significantly lower background level of reads compared to well-watered leaf tissue. This enrichment resulted in a higher abundance of reads in open peaks, potentially indicating a major difference in chromatin state in the nucleus of desiccated cells.

We hypothesize several factors that may contribute to the low background signal and robust peaks observed under desiccation. The first is that all nuclei in the desiccated plants are more condensed either due to crosslinking or the presence of additional factors that prevent the Tn5 from sometimes cutting. The more parsimonious explanation is that the desiccated nuclei are in a more stable state with fewer cells in the sample actively undergoing transient processes that open the DNA, such as transcription or replication. Under desiccation, different cell types must arrest their normal developmental or metabolic functions and enact a series of highly coordinated responses to successfully prepare for anhydrobiosis and subsequent rehydration. In turn, this would ‘synchronize’ cells and produce the observed tight coordination between expression dynamics and chromatin architecture that is typically only found in single cell data (Farmer et al., 2021). Drying of this magnitude is experienced by all cells and risky processes such as growth and photosynthesis during the stresses of anhydrobiosis would be detrimental and preferentially avoided. Thus, a clear, tightly regulated signal from all cells in the desiccated samples is achieved. This is supported by the massive transcriptional reprogramming we observed and the clear enrichment of GO terms related to photoprotective and anhydrobiosis processes with comparatively few background pathways. This pattern of clearer peaks and more accessible chromatin under desiccation contrasts what has been observed under drought and salinity (Raxwal et al., 2020), but is similar to cold stress (Zeng et al., 2019).

### Evolutionary dynamics of photoprotective strategies under desiccation

*O. thomaeum* and *E. nindensis* are found within the Chloridoideae subfamily of grasses, a group of stress tolerant C4 species with superior drought, heat, and salinity tolerance (Marcum, 1999; Peterson et al., 2001). Evolutionary access to existing resilience traits within this subfamily likely enabled the independent evolution of desiccation tolerance in *O. thomaeum* and *E. nindensis* (Pardo and VanBuren, 2021). Using detailed comparative genomics approaches with syntenic orthologs, we identified a core set of genes and *cis*-regulatory regions that are induced during desiccation in *O. thomaeum* and *E. nindensis*. Integration of ATACseq and RNAseq data allowed us to filter out spurious associations, and induced genes include many with previously described roles in desiccation. Among the stress responsive *cis*-elements, we identified numerous binding motifs for the seed desiccation transcription factor ABI5, supporting the hypothesis that vegetative desiccaion tolerance evolved from rewiring existing seed pathways (Oliver et al., 2000, 2005). This hypothesis is contentious, and previous work in the monocot *Xerophyta humilis* failed to find evidence linking desiccation processes to the canonical seed development transcription factor network of LEC1, ABI3, ABI5 and others (Lyall et al., 2020). Our results provide a direct link between expression dynamics and regions of accessible chromatin with seed related regulatory motifs, but this association needs to be further tested using transcription factor binding data.

Species-specific changes in chromatin architecture and expression dynamics during desiccation reflect the distinct strategies that *E. nindensis* and *O. thomaeum* utilize to mitigate photooxidative damage. We identified a cluster of desiccation associated GO terms related to chlorophyll processes, UV-A responses, photoprotection, and pigment metabolism that was unique to *O. thomaeum*. These orchestrated responses in *O. thomaeum* reflect homoiochlorophyly, or the strategy to protect chlorophyll and photosynthesis related macromolecules during desiccation. Consistent with this, *O. thomaeum* has more early light induced proteins than *E. nindensis* (when accounting for ploidy) and all of the ELIPs have massive shifts in chromatin accessibility and high expression under desiccation. By comparison, only two ELIPs in *E. nindensis* have associated open chromatin regions, and few are differentially expressed under desiccation. Instead, ELIPs in *E. nindensis* have high expression during rehydration and likely function in protecting leaves as they resynthesize and repair their photosynthetic apparatus (Pardo et al., 2020).

We identified a *cis*-regulatory motif associated with the drought transcripton factor DREB upstream of each ELIP in *O. thomaeum*. This motif is missing from the Arabidopsis ELIP orthologs and may represent a regulatory neofunctionalization to induce ELIP accumulation under desiccation. Nearly all plants have retained desiccation tolerance in their seeds and/or pollen, so the genes and regulatory elements needed to protect cells from anhydrobiosis are already in place. Tolerance may have evolved in resurrection plants through simply shifting the timing and cell specificity of existing desiccation pathways through cis-regulatory elements, and our findings in ELIPs may be evidence of this process in action.

The GO-terms we found imply desiccated plants were under stress responding to desiccation, and a general down-regulation of photosynthesis and carbon fixation in desiccated plants compared to the well-watered. Both species also have an increase in genes associated with production of vitamin B6 under desiccated conditions. While vitamin B6 is associated with scavenging reactive oxygen it is unclear if B6 levels are higher in desiccated tissues or what function these compounds are performing. Desiccation in *O. thomaeum* also resulted in several GO-terms implying responses to high light intensity, and a preservation of light-harvesting and dealing with cellular stresses of excess light, whereas in *E. nindensis* there were more terms pertaining to catabolism of branched-chain amino-acids, endocytic recycling, and autophagy. Additionally, while both species had a down regulation of photosystem I, *E. nindensis* also had a down-regulation of photosystem II. These differences highlight the fact that while both species respond to the loss of water similar to many other plants, they also have processes that seem specific to each of them.

Genes with higher differential expression in drought typically have a nearby chromatin region that is also differentially open during desiccation. However, there are some differentially expressed genes that have opposing open chromatin regions and these genes are enriched in house-keeping processes such as polysaccharide biosynthesis, ATP mediated transport, rRNA biosynthesis, and nuclear envelope organization (Supplementary table 8). This implies coordinated gene expression that is contrary to the simplistic thinking of “more open chromatin results in more transcript abundance”. This observation could be attributed to repressors binding in or near these open sites, post-transcriptional control of mRNA levels, or priming of gene expression for returning to a hydrated state. A time course of expression through the desiccation process could help identify what is happening in these cases.

## Methods

### Plants, samples, and growth conditions

*Oropetium thomaeum* and *Eragrostis nindensis* plants were grown from seed for ∼60 days for each experiment in a growth chamber with 28 °C (day)/22 °C (night) and 12 hr light/12 hr dark cycle. Desiccated samples were collected from plants where water was withheld for 10 days, causing the leaf relative water content to drop below 10%. Each replicate consisted of 3 mature plants grown in the same pot, and two replicates were collected for well-watered and desiccated time points. Two grams of leaf tissue was collected from each plant and frozen in liquid nitrogen immediately. Tissues were ground into coarse powder and aliquoted into multiple 1.5 mL tubes with ∼100 mg per tube. The samples were then subjected to nuclei isolation for ATAC-seq, or RNA extraction for RNA-seq.

### RNA-seq library construction

RNA was extracted using Zymo research Direct-ziol RNA miniprep kit according to the manufacturer’s protocol with on-column DNase digestion. RNA yield was quantified using a Qubit RNA BR kit and RNA integrity was quantified by gel electrophoresis. One microgram of RNA was used to construct the RNA-seq libraries using Illumina TruSeq stranded mRNA kit, according to the manufacturer’s protocol. The multiplexed libraries were sequenced at Michigan State University RTSF Genomics Core with HiSeq4000 150 bp paired end mode for *O. thomaeum* and 100 bp paired end mode for *E. nindensis*.

### ATAC-seq library construction

Crude nuclei extracts were prepared using the protocol reported in Liu et al. (2017) (Lu et al., 2017). Nuclei were stained with 4,6-Diamidino-2-phenylindole (DAPI) and counted using a Nikon Eclipse Ni-Upright microscope with 40X differential interference contrast objective lens. 50,000 nuclei were subjected to 2 μL of Tn5 enzyme digestion for 30 min at 37 °C. DNAseq libraries were constructed from the resulting digested nucleosomes using the Illumina Nextera XT kit, according to the manufacturer’s protocol. The multiplexed libraries were sequenced at Michigan State University RTSF Genomics Core with HiSeq4000 150 bp paired end mode.

### RNAseq analysis

Paired-end raw reads were trimmed using Trimmomatic (v0.33) (Bolger et al., 2014) to remove adapters and low-quality bases. Reads were pseudo-aligned to the *O. thomaeum* v2.1, or *E. nindensis* v2.1 transcriptome using Kallisto (0.46.0) (Bray et al., 2016). Differential expressed genes (q-value < 0.05) between well-watered and drought conditions were identified using Sleuth (Pimentel et al., 2017).

### ATAC-seq analysis

Paired-end raw reads were trimmed using Trim_galore (v0.6.6) to remove Nextera adapters and low-quality bases (Krueger, 2018). Filtered reads were then aligned to *O. thomaeum* v2.1, or *E. nindensis* v2.1 genome using Bowtie2 with dovetailing reads enabled and the resulting SAM files were converted to BAM format and sorted based on query name using Picard Tools (v2.18.1) (Institute, 2016). Mapped reads were manually visualized using the Integrative Genomics Viewer (Robinson et al., 2011), to determine uniformity of coverage. A blackout list of genomic regions with abnormal read coverage was generated by investigating genes near large regions with greater than 100x coverage relative to other parts of the chromosome. Regions with both exceptionally high coverage and genes associated with the chloroplast were considered chloroplast contamination in the build of the genome. A total of 25 regions consisting of about 1.5 Mbp of the ∼236 Mbp *O. thomaeum* genome, and 772 regions consisting of about 3.5 Mbp of the ∼986 Mbp *E. nindensis* genome, were added to the blackout lists.

Peaks associated with accessible chromatin regions (ACR) were called using Genrich (Gaspar) with options specific for ATACseq. The resulting narrowPeak files and the read files (bam format) were used in the R library DiffBind (Stark et al., 2011; Ross-Innes et al., 2012) to determine differentially open regions. Running the analysis with bFullLibrarySize=FALSE both edgeR (Robinson et al., 2010) and DESeq2 (Love et al., 2014) produced similar outcomes in *O. thomaeum* so the more conservative edgeR results were used for downstream analyses. In *E. nindensis* the results were more lopsided so the better distributed DESeq2 results were used. (Supplemental tables)

### Integrating ATACseq and RNAseq datasets

A genome-wide assessment of ACR distribution was performed using deepTools (Ramírez et al., 2016). ACRs were classified into different groups based on their proximity to genes and comparisons of peak openness and differential gene expression were performed. These correlations were used as input for gene ontology and cis-element enrichment analyses.

We identified genes near ACRs using bedTools version 2.29.2 (Quinlan, 2014). ACRs were grouped into different categories based on their distance to genic elements including: 10000 to 5001, 5000 to 2001, 2000 to 1001, 1000 to 501, 500 to 0 bp upstream (5’) of the transcription start site (TSS), overlapping with the gene, and 0 to 1000bp downstream (3’) of the transcription termination site (TTS). Based on the distribution and correlation of chromatin openness and gene expression patterns, we chose the range of 0-3000 bp upstream of gene TSSs as the cutoff of associating ACRs with specific genes for further analysis. The *O. thomaeum* and *E. nindensis* genomes are relatively compact compared to other grass species, and putative cis-regulatory regions are in close proximity to genes.

Genes were grouped by their differential expression patterns and ACR openness and gene ontology term (GO-term) enrichment analyses were performed using the R library topGO (Alexa and Rahnenführer, 2009). Six different sets of genes from each species were used as inputs for GO term enrichment including up and down-regulated genes under desiccation with no comparison to ACRs, and the four categories of gene expression and ACR openness association including: (1) upregulated genes near more open ACRs under desiccation, (2) less open ACRs and downregulated expression (3) less open ACRs and upregulated expression or (4) upregulated expression less open ACRs.

Putative *cis*-regulatory motifs within ACRs were identified using Multiple Em for Motif Elicitation (MEME) (Bailey et al., 2009; Bailey, 2020). Motifs associated with ACRs within 3 Kbp upstream of differentially expressed genes were identified and the Simple Enrichment Analysis (SEA) algorithm was used to identify enriched motifs in our data and associate these motifs with known transcription factor binding sites (Bailey and Grant, 2021). Sensitive Through Rapid Em Motif Elicitation (STREME) was used for denovo motif identification (Bailey, 2020) for enriched motifs with no homology to known transcription factor binding sites.

### Comparative genomic analyses

Comparative genomic analyses were conducted between *O. thomaeum* and *E. nindensis* to identify shared and species-specific sets of differentially expressed genes and ACRs dynamics. Syntenic orthologs were identified between the two closely related grasses using the python version of MCScan (https://github.com/tanghaibao/jcvi/wiki/MCscan-(Python-version)) (Wang et al., 2012). The chromosome scale *O. thomaeum* genome was used as an anchor and the two sets of homeologous genes in the *E. nindensis* genome were mapped to the single corresponding syntenic ortholog in *O. thomaeum*. Genes in each species were then classified as syntenic or non-syntenic and these two designations were incorporated with differential expression analyses, ACR dynamics, and enriched motif analyses. Enriched GO terms and putative *cis*-element motifs were identified for syntenic orthologs with conserved and species-specific expression patterns as described above.

## Data availability

The raw RNAseq and ATACseq data are available from the National Center for Biotechnology Information (NCBI) Short Read Archive. Raw data for this project can be found under BioProject accession no. PRJNA807505.

## Acknowledgments

This work is supported by NSF Grant MCB□1817347 (to R.V.).

