## Supplemental Figures/Tables for "Regulatory dynamics distinguishing desiccation tolerance strategies within resurrection grasses"

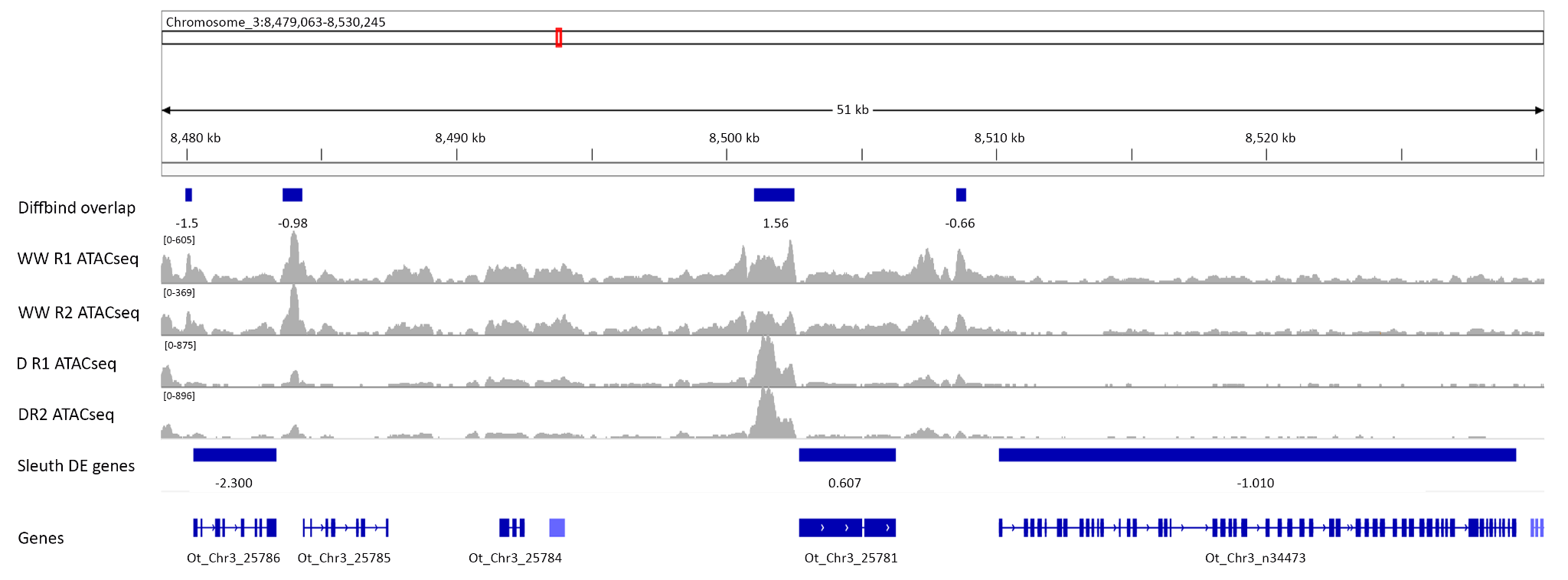

**Fig S1: Genome Browser view of ATAC-seq and RNA-seq data near a region with differentially expressed genes in *Oropetium thomaeum*.**

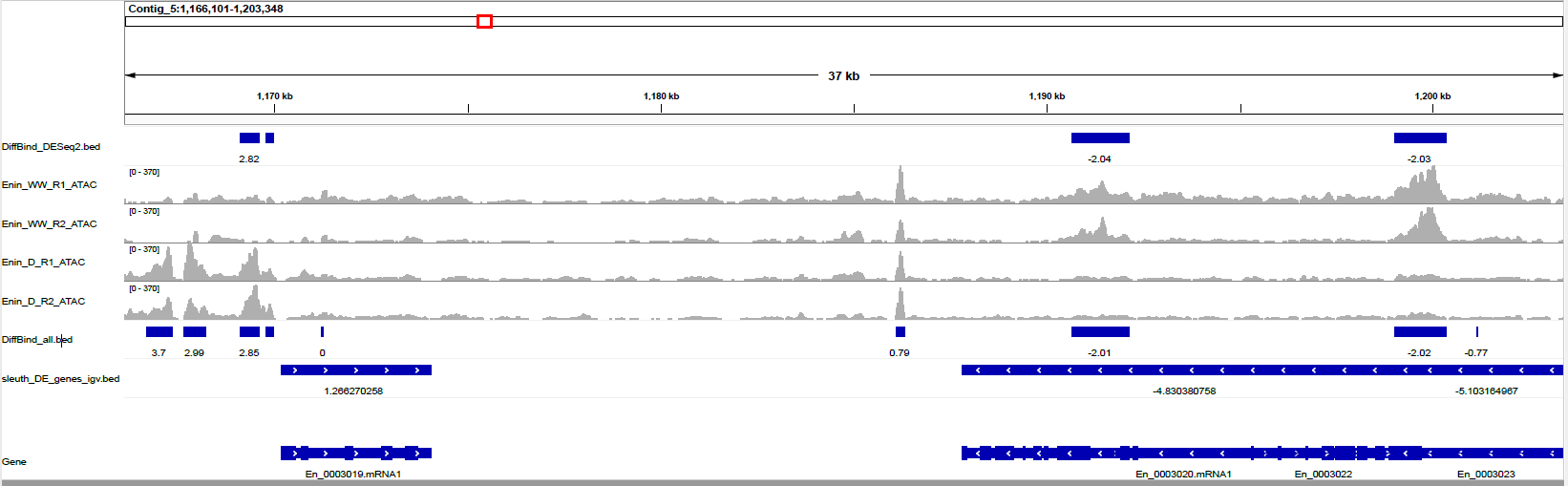

**Fig S2: Genome Browser view of ATAC-seq and RNA-seq data near a region with differentially expressed genes in *Eragrostis nindensis*.**

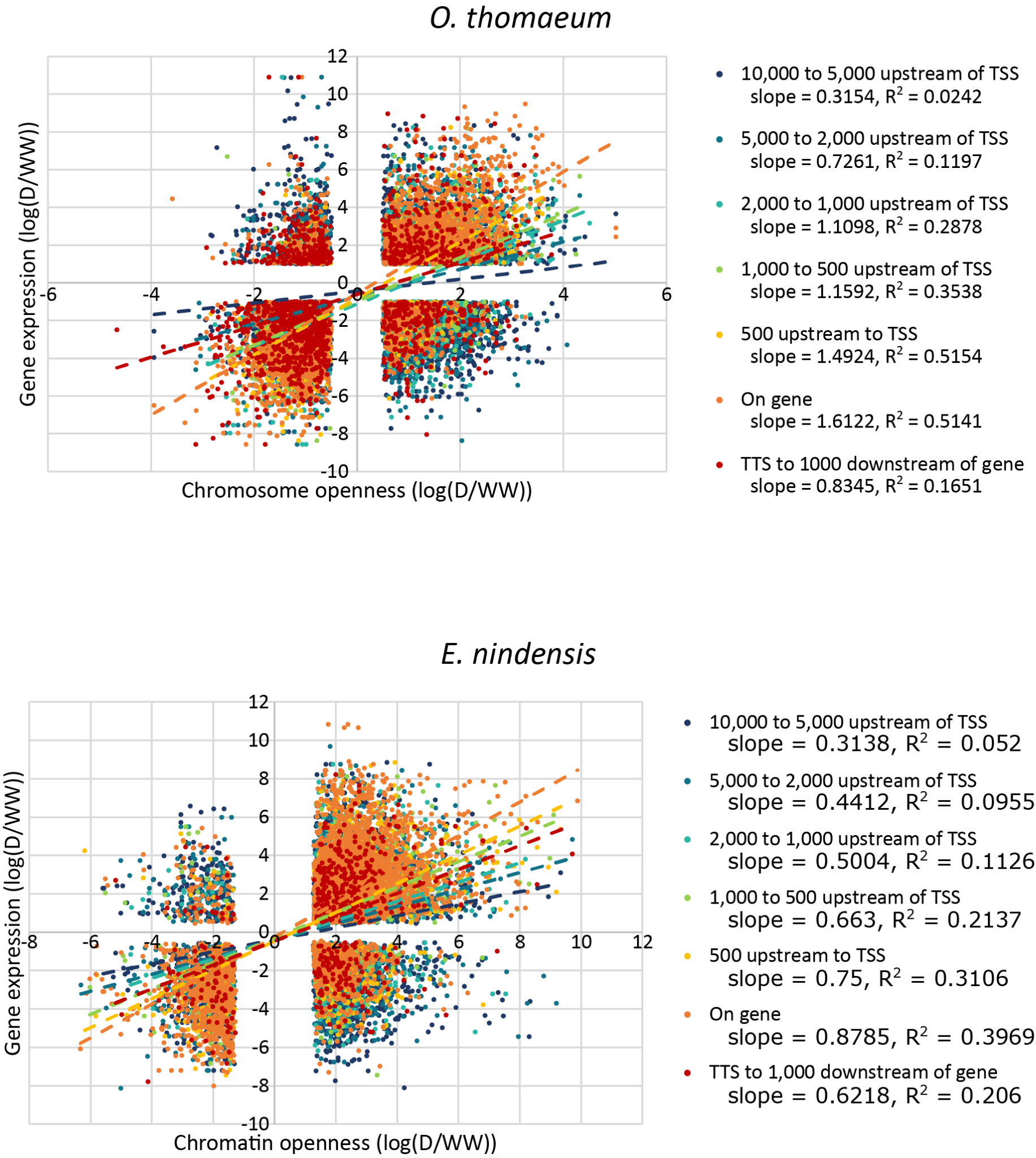

**Fig S3: Correlation between differential gene expression and nearby chromatin openness in (a) *O. thomaeum* and (b) *E. nindensis*.** The differentially expressed genes in desiccation with differentially open chromatin within denoted regions are plotted where genes in the upper right hand quadrant represent genes that are more open with higher expression under desiccation. Genes in the lower left hand quadrant have lower expression and less chromatin openness under desiccation.

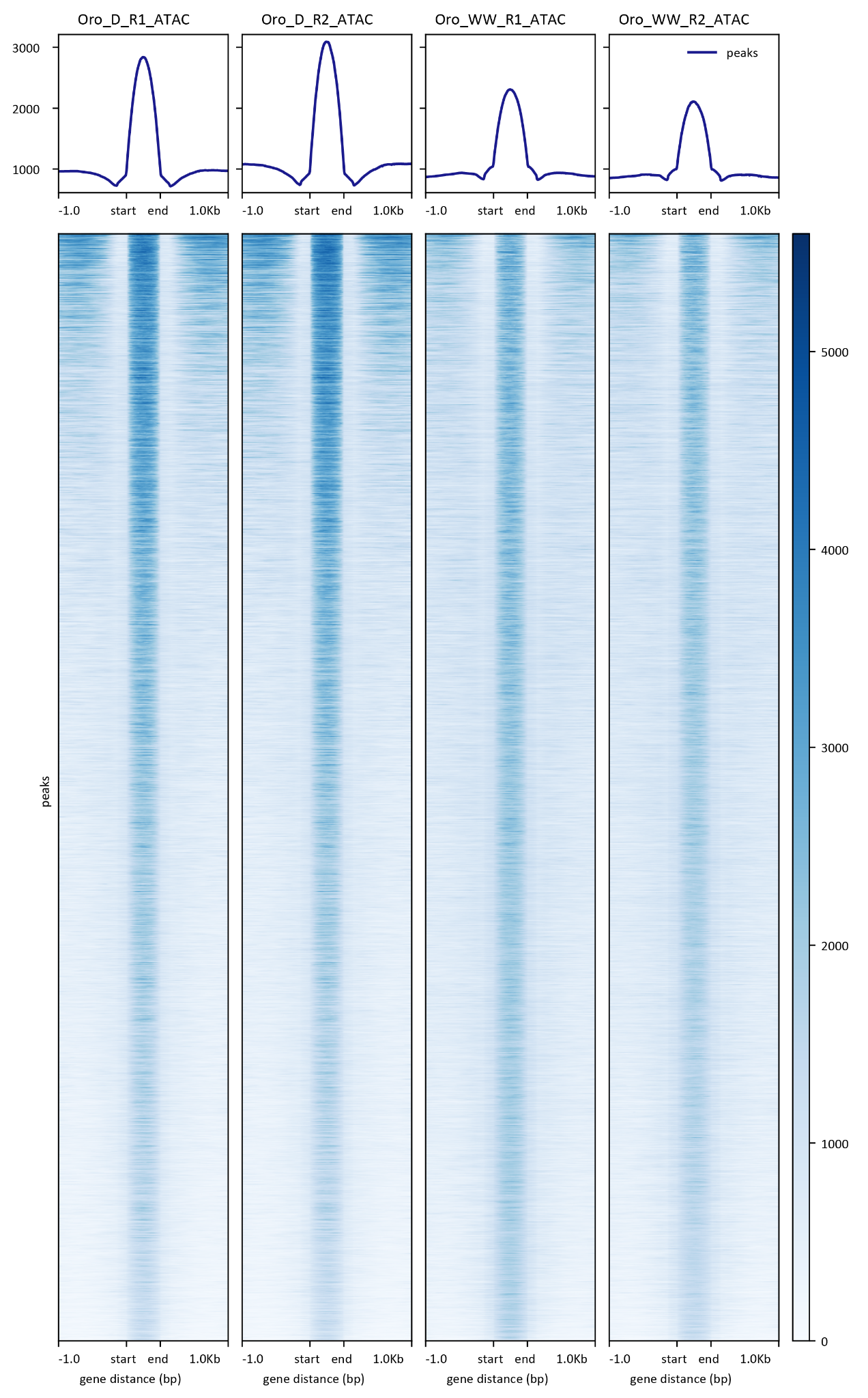

**Fig S4: Scaled plot of open regions and surrounding chromatin area in *O. thomaeum*.**

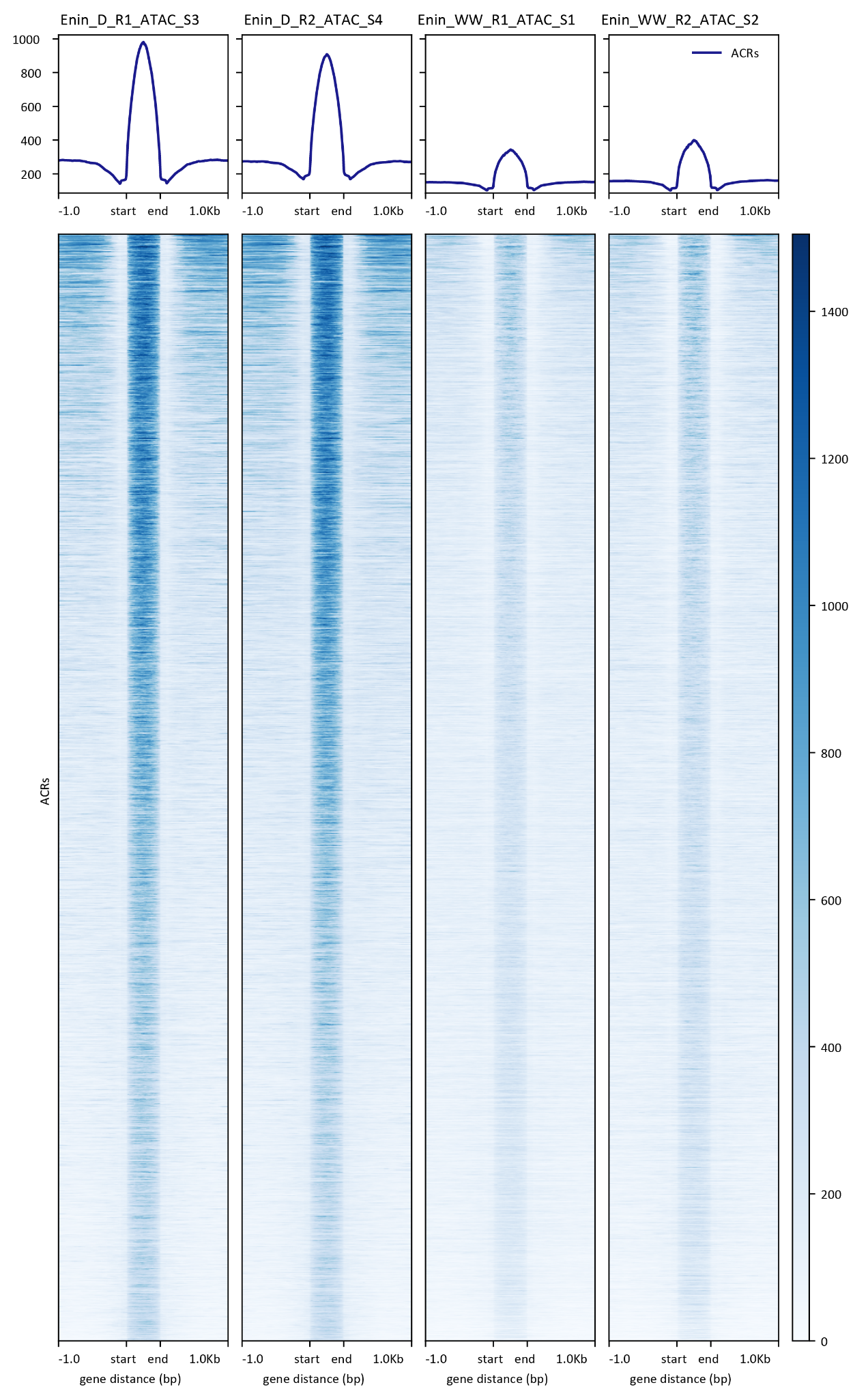

**Fig S5: Scaled plot of open regions and surrounding chromatin area *E. nindensis*.**

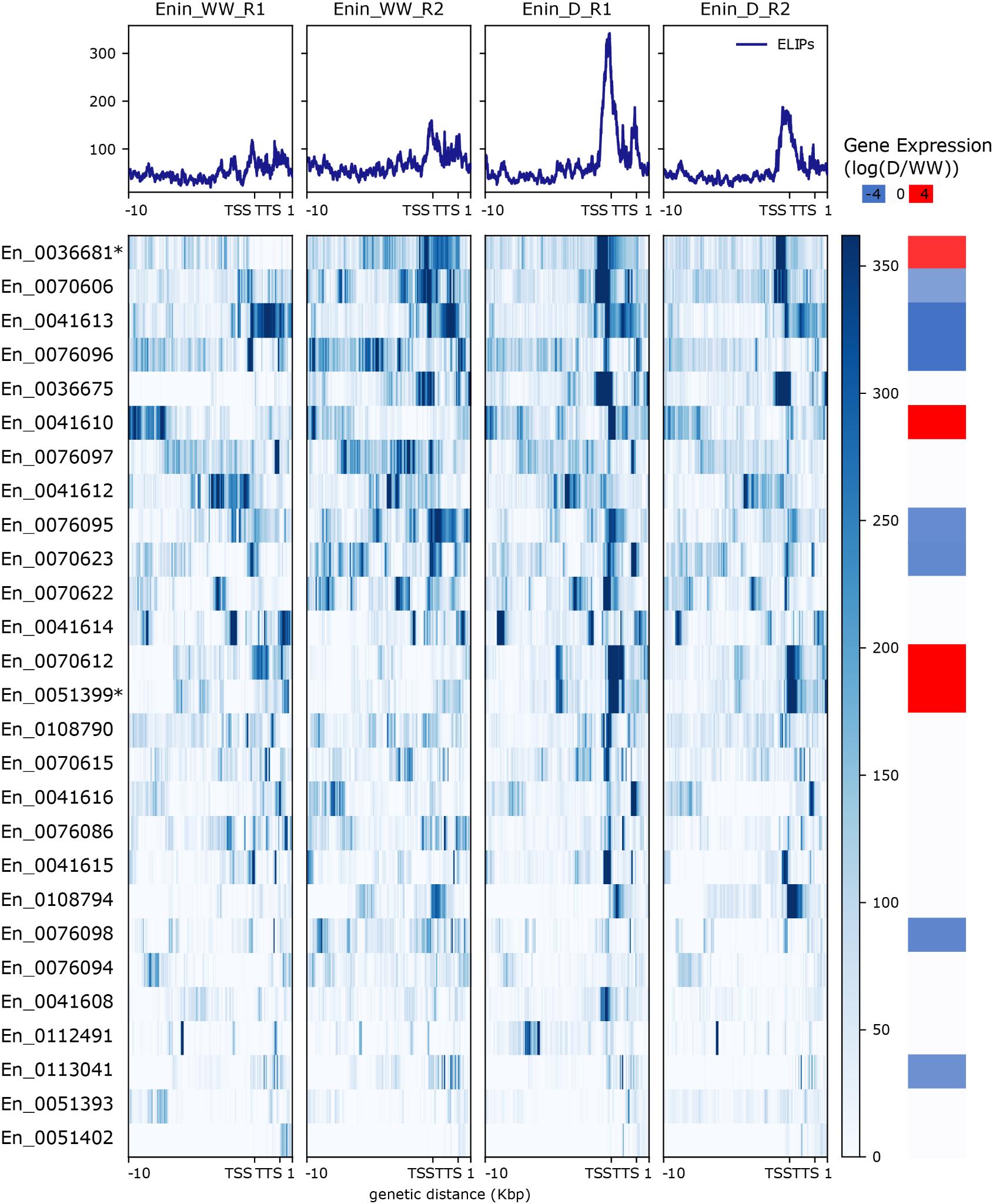

**Fig S6:** **Regulatory dynamics of ELIPs in *E. nindensis.*** Chromatin architecture and expression dynamics of ELIPs in well-watered and desiccated *E. nindensis* samples. The mean mapped read depth of ATACseq reads (in RPKM) is plotted for 10 kb upstream to 1 kb downstream regions for each of the ELIPs in the *O. thomaeum* genome. Log2 transformed RNA expression (in TPM) for each ELIP is shown on the right under well-watered and desiccated conditions.

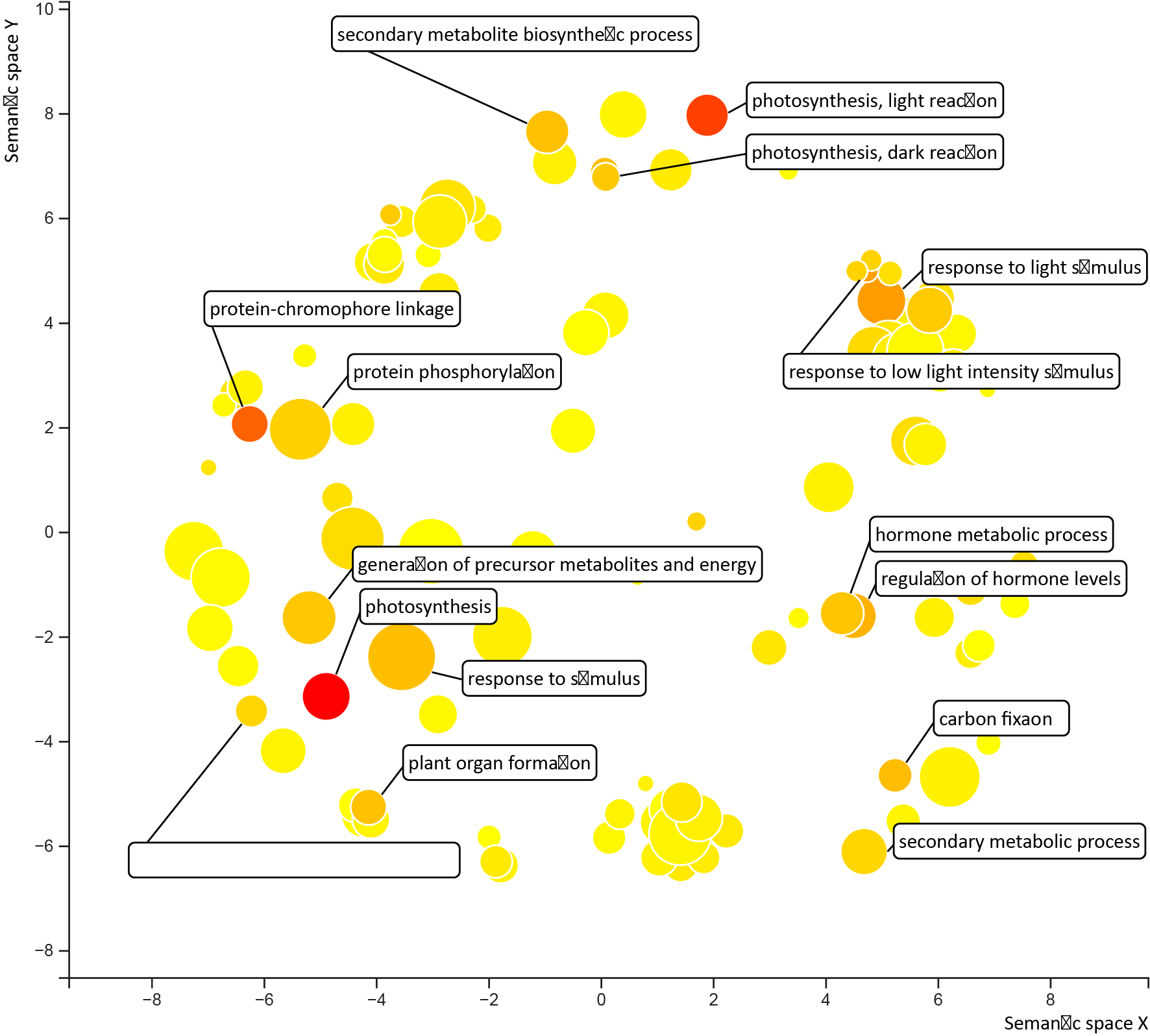

**Fig S7: Enriched GO terms for genes that have lower expression and less open ACRs under desiccation are plotted for *O. thomaeum*.** GO terms are transformed using Multidimensional Scaling to reduce dimensionality and terms are grouped by semantic similarities. GO terms with previously characterized roles in desiccation and photoprotection responses are highlighted. The color of the circles represent significance and size represents the number of genes in that group.

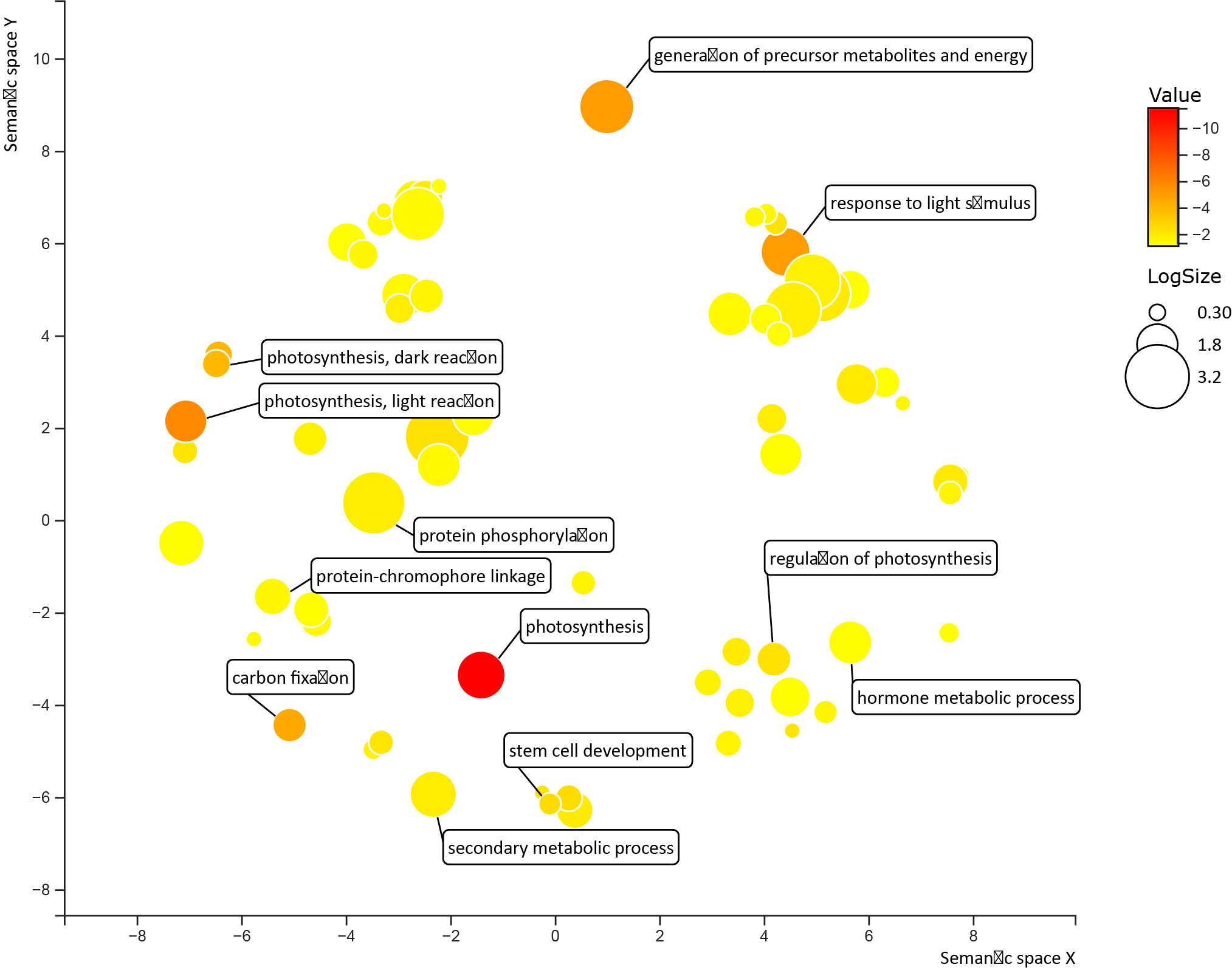

**Fig S8: Enriched GO terms for genes that have lower expression and less open ACRs under desiccation are plotted for *E. nindensis*.** GO terms are transformed using Multidimensional Scaling to reduce dimensionality and terms are grouped by semantic similarities. GO terms with previously characterized roles in desiccation and photoprotection responses are highlighted. The color of the circles represent significance and size represents the number of genes in that group.

**Supplemental Tables**

Supplemental Table 1: Differential gene expression in *O. thomaeum*

Supplemental Table 2: Differential gene expression in *E. nindensis*

Supplemental Table 3: Differential chromatin openness in *O. thomaeum*

Supplemental Table 4: Differential chromatin accessibility in *E. nindensis*

Supplemental Table 5: Syntenic differential expression in *O. thomaeum* and *E. nindensis*

Supplemental Table 6: Syntenic chromatin accessibility in *O. thomaeum* and *E. nindensis*

Supplemental Table 7: Gene Ontology Heatmap

Supplemental Table 8: Gene Ontology for different sets of genes and chromatin openness

Supplemental Table 9: Gene Ontology for different sets of syntenic genes
